# Proteomics of Transglutaminase 2 polyaminated proteins

**DOI:** 10.1101/2025.05.28.656543

**Authors:** Don Benjamin, Danilo Ritz, Sunil Shetty, Sujin Park, Michael N. Hall

## Abstract

Post-translational modification of proteins by polyamines (putrescine, spermine and spermidine) is poorly studied due to technical challenges in identification. Polyamination is mainly performed by transglutaminases via transamidation of a polyamine to an acceptor glutamine-residue on the target protein. We performed polyamination reactions on whole cell lysates using 2 different polyamine probes (Biotin-pentylamine and DNP-pentylamine) and identified modified peptides by mass spectrometry. 51 protein targets modified on 66 different sites were identified, with an overlap between hits tagged with the different probes. Many of the targets are involved in translation, cytoskeletal organization or have roles in cancer.

## Introduction

Polyamines are an essential metabolite for cellular health, DNA replication, gene expression, translation and proliferation (1). The major polyamine species are putrescine, spermine and spermidine, which are produced as an off-shoot of arginine metabolism. Cadaverine, a product of bacterial metabolism in the gut, is also found in the serum. Despite their wide-ranging roles, only a few specific roles are known for the polyamines, and most of its effects in the cell are attributed to their net positive charge. The best characterized specific role is for spermidine which is a precursor for the formation of hypusine on eIF5A (2). Nonetheless, post-translational modification of proteins by polyamination at glutamine residues has long been known (3), and in a few cases the consequence of this modification has been elucidated in some detail. The Bordetella dermonecrotizing toxin polyaminates mammalian Rho at Q63 (and analogous sites on Rac1 and Cdc42) and elicits infectious toxicity (4). Mostly however, protein polyamination is performed by endogenous transglutaminases (TGM), a large family of proteins of which the best studied is Transglutaminase 2 (TGM2). RhoA polyaminated by TGM2 has been shown to be constitutively activated (5, 6). TGM2 polyaminated phospholipase A2 is also hyperactivated (7). TGM2 polyamination stabilizes tubulin (8) while polyaminated lipocalin 2 is rapidly cleared (9). Recently, the 4EBP proteins were shown to be polyaminated by TGM2 under hypoxic conditions to favor selective mRNA translation (10). TGM2 also polyaminates BAF250a to modulate expression of selective genes (11). Thus, there are examples where polyamines exert an effect not merely due to mass action as abundant positively charged molecules (they are typically present at millimolar concentration), but in a highly specific manner.

Proteomics of polyaminated proteins has lagged behind characterization of other PTMs due to the incompatibility of the polyaminated adduct to standard MS methodology, resulting in its loss during sample purification, or destruction during data acquisition. The most recent proteomic study used an anti-spermine antibody to pull down in vitro polyaminated proteins from cell lysates and identified 254 polyaminated sites from 233 proteins (12). Apart from this, there is an absence of large-scale surveys and most described examples have come via a bottom-up approach from investigation of individual proteins. In most of these studies, except for rare exceptions, the identity of the modified glutamine is not known.

We developed a cell line that depends exclusively on TGM2 for polyamination. Using 2 differently tagged polyamine probes, we identified 51 target proteins (66 sites). There is significant overlap between the sites labelled by the different probes, giving confidence to the data. Some of the hits have been previously reported but we also report a significant number of novel hits. In particular we could identify the glutamine acceptor sites for all of the hits, something missing in most previous reports. Among the targets are proteins involved in translation, cytoskeletal organization and cancer.

## Results

### Characterization of CB1 as exclusively dependent on TGM2 for polyamination

Polyamination is a transamidation reaction where a polyamine is covalently attached by its amine group to the *γ*-carboxamide of a recipient glutamine on a protein (Fig 1A). This is performed by TGMs that attach not just polyamines but any primary amine to an acceptor glutamine (for example serotonylation or dopaminylation). TGMs are multifunctional enzymes (13). There are 7 TGMs and an eighth protein, FactorXIIIA, in mammals that perform protein polyamination. The most widely expressed TGM isoform, and the best studied, is TGM2.

**Figure 1.**
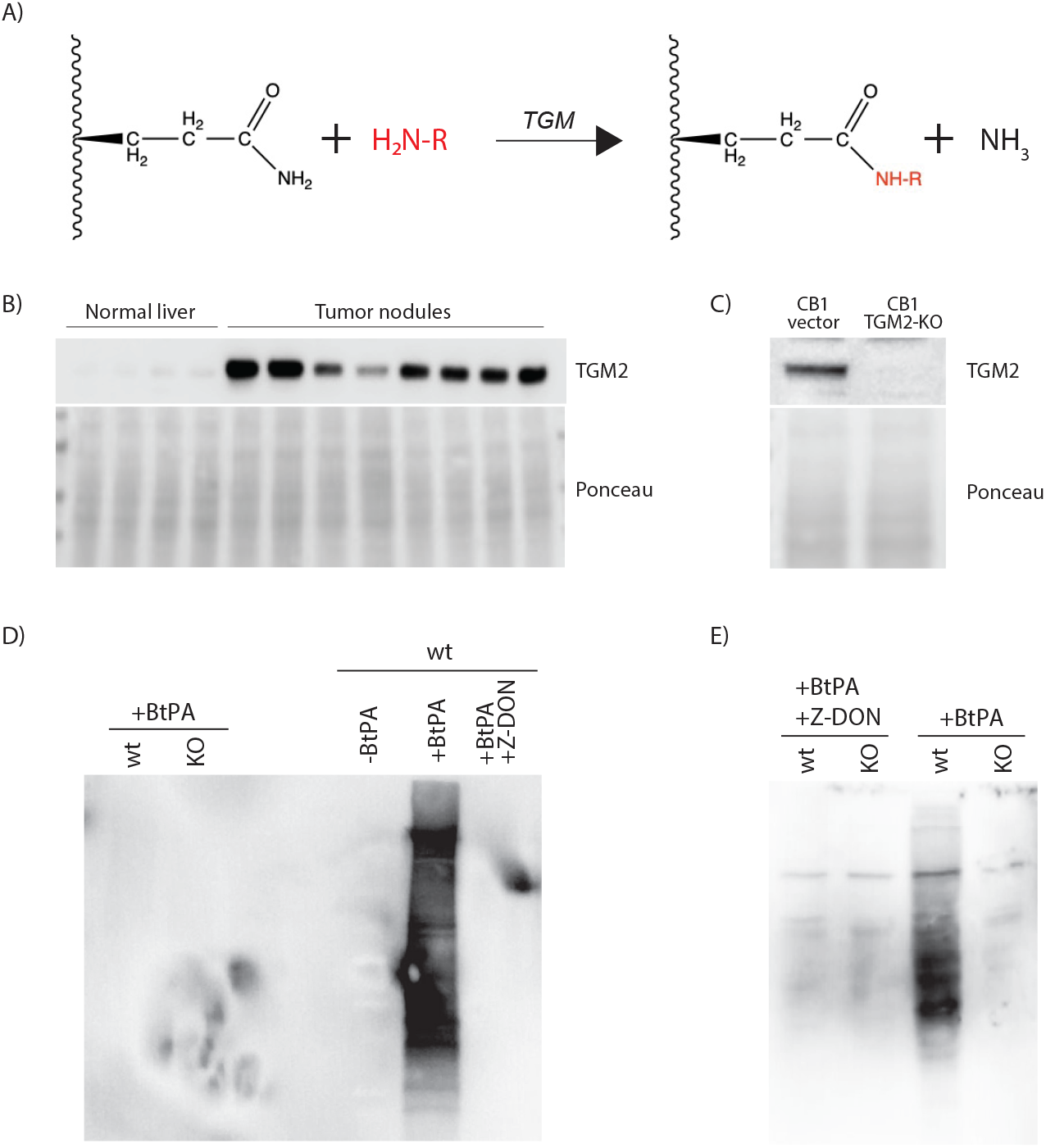
(A) Transamidation of an acceptor glutamine on a target protein by a transglutaminase. If the moiety on the primary amine is a polyamine, this will result in polyamination of the target protein. (B) Over-expression of TGM2 in liver tumor nodules compared to normal liver tissue. (C) Crispr knock-out of TGM2 in CB1 cells. (D) In vitro polyamination of CB1 cell lysate with biotin-pentylamine (Bt-PA). (E) Incorporation of Bt-PA by CB1 cells and polyamination of proteins in vivo. Blots were probed with HRP conjugated streptavidin (SA-HRP).

We identified TGM2 as being over-expressed in the CB1 cell line that was derived from mouse hepatocellular carcinoma tumors (Fig 1B) (14). TGM2 was knocked out using Crispr (Fig 1C), and lysates from parental and TGM2-KO cells were assayed for TGM2 dependent polyamination activity using biotin-tagged pentylamine (Bt-PA). Bt-PA has a 5-carbon long polyamine moiety similar to cadaverine (Fig 2A). Bt-PA was incorporated into endogenous proteins in wild-type CB1 lysate but not in corresponding lysate from TGM2-KO cells (Fig 1D). Bt-PA incorporation is completely blocked by the TGM2-specific inhibitor Z-DON. Similar observations are seen with in vivo incorporation and labelling with Bt-CAD (Fig 1E).

**Figure 2.**
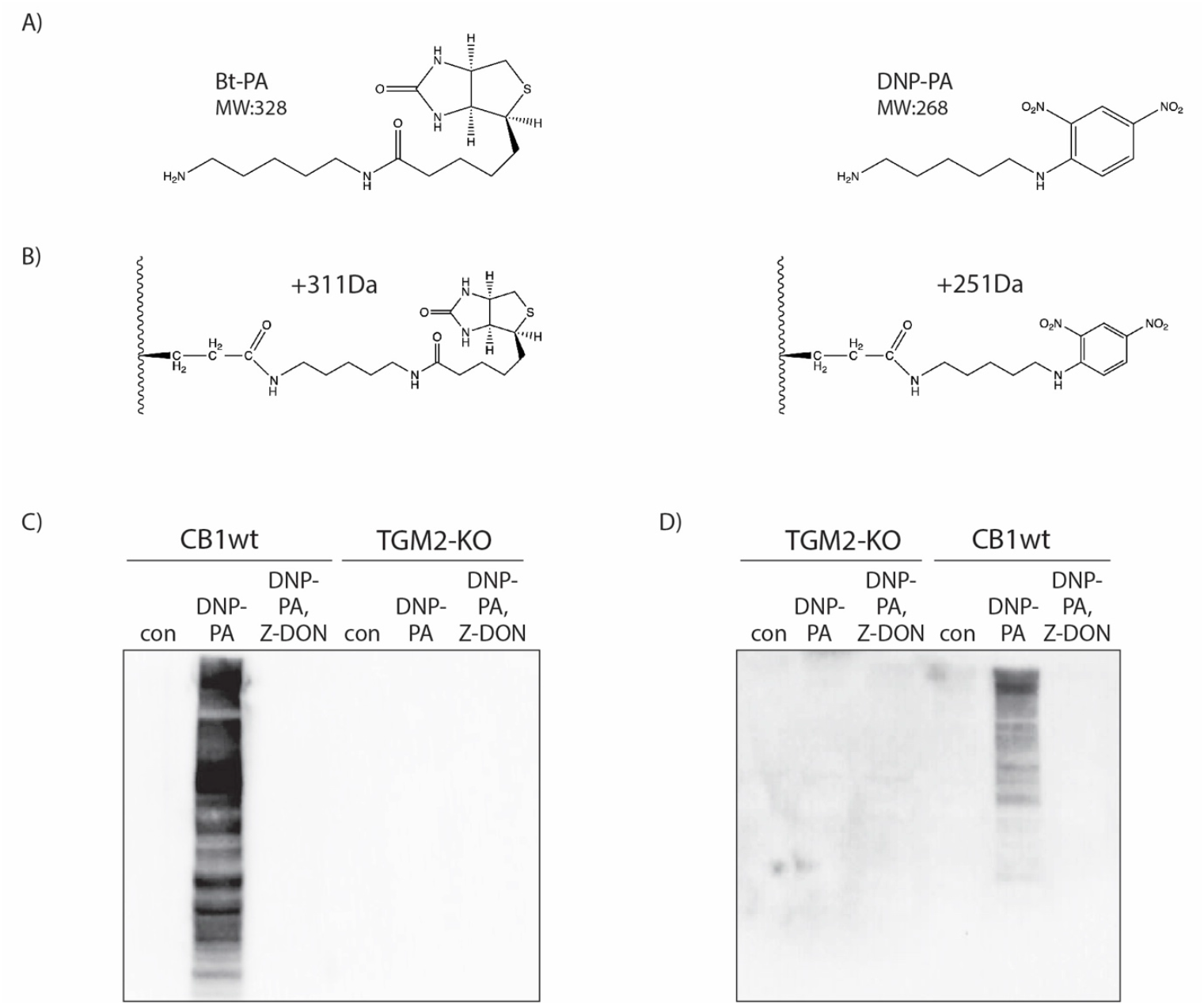
(A) Structures of biotin-pentylamine (Bt-PA) and DNP-pentylamine (DNP-PA). (B) PTM of glutamine with Bt-PA or DNP-PA and the predicted mass shift. (C) In vitro polyamination of CB1 cell lysate with DNP-PA. (D) Incorporation of DNP-PA by CB1 cells and polyamination of proteins in vivo. Blots were probed with anti-DNP antibody.

Taken together, the results demonstrate that CB1 relies solely on TGM2 for protein polyamination and is a suitable choice as a model system to study TGM2-dependent polyamination.

### In vitro polyamination and enrichment of labelled proteins

As polyamination activity is provided solely by TGM2 in CB1, we could directly use whole cell lysates to perform labelling with tagged polyamines using endogenous TGM2 present in the lysate. Two different polyamine tags were used for labelling: Bt-PA and DNP-PA (Fig 2A). The different sets of reagents and downstream processing post-labelling for each label also helps to rule out systematic error in peptide enrichment. Additionally, the different molecular weights of the tags will be important during data analysis where we hope to detect the predicted mass shifts in the spectra corresponding to the tag employed (Fig 2B).

We performed labelling with DNP-PA on CB1 using the same conditions as for Bt-PA and confirmed TGM2-specific labelling with DNP-PA both in vitro and in vivo (Fig 2C).

### MS identification of peptides enriched for polyaminated tags

For MS identification of polyaminated peptides, the following work-flow was set up (Fig 3A). In vitro polyamination reactions of CB1 lysates were performed in triplicate either (i) without tagged-PA, (ii) with tagged-PA, and (iii) with tagged-PA and Z-DON. This block of reactions was repeated twice making for 3 biological replicates in total. Under this schema, we expect to detect modified peptides only under condition (ii) with conditions (i) and (iii) serving as internal controls. We also performed triplicate reactions using Bt-PA with TGM2-KO lysate as a further negative control of our methodology.

**Figure 3.**
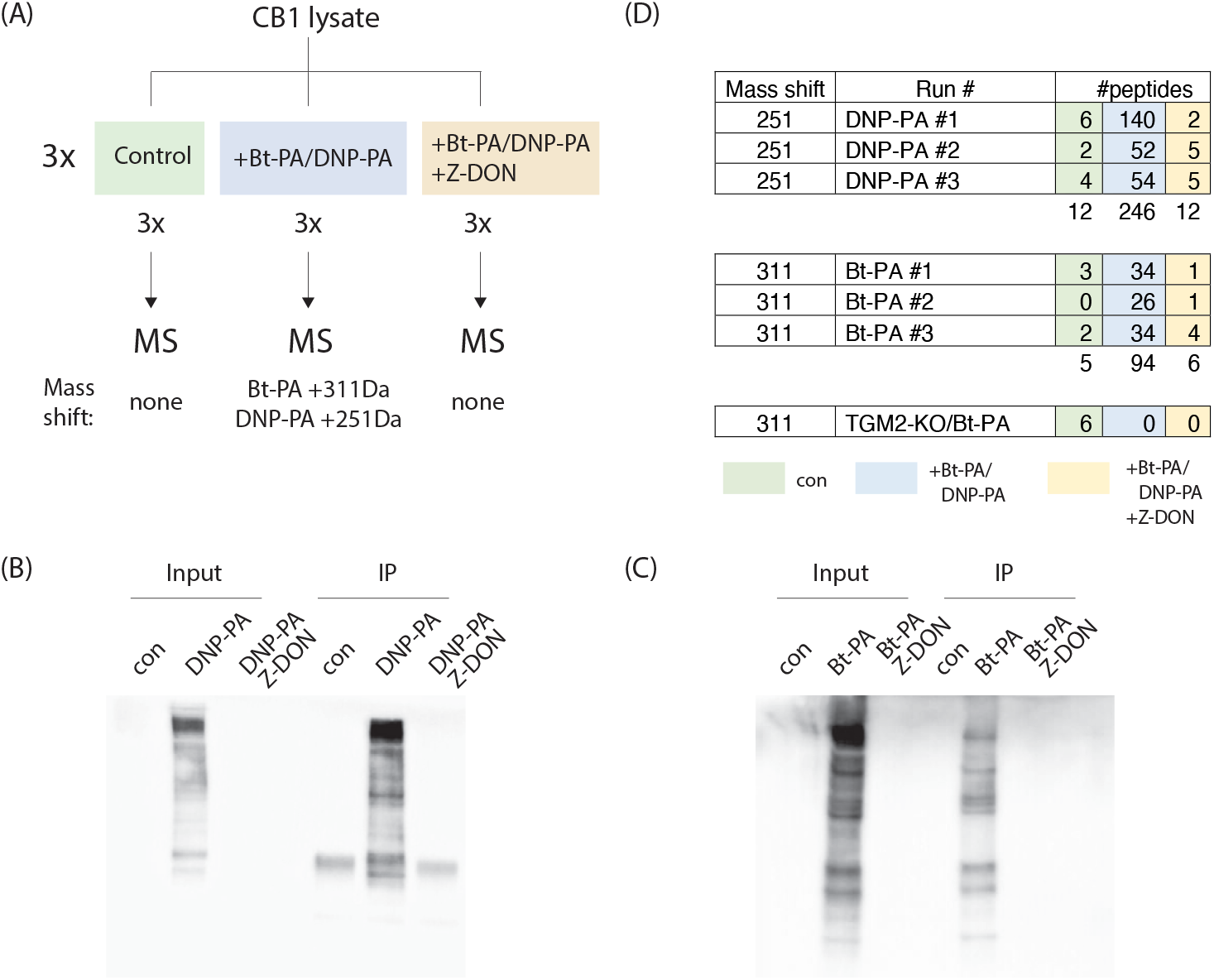
(A) Work-flow for detection of TGM2 polyaminated targets. Peptides bearing expected mass shifts are only expected in the condition with label added and no TGM2 inhibition (blue). (B, C) Examples of quality control performed for each experiment to show labelling and enrichment before sending IP samples for MS. Blots were probed with anti-DNP antibody or SA-HRP. (D) Tabulation of number of peptides detected for the workflow presented in (A), and for a control experiment using TGM2-KO lysate.

Polyaminated proteins were enriched by immunoprecipitation with agarose bound anti-DNP and anti-biotin antibody beads (Fig 3B). Enrichment of Bt-PA tagged proteins by streptavidin beads gave poor MS results regardless whether the peptides were liberated by boiling or on-bead digestion, leading us to switch to anti-biotin antibody beads for immunoprecipitation.

Antibody-bound proteins were subjected to on-bead trypsin digestion and eluted peptides processed as described in Methods. Samples were then analyzed by LC-MS/MS.

### Identification of peptides with expected mass shifts

Acquired raw-files were analyzed by open search to identify delta mass of either +251.0906 Da or +311.16675 Da corresponding to which tag was employed (DNP-PA and Bt-PA respectively). The majority of peptides shifted by the predicted mass values were detected in the expected condition (+tagged-PA), with significantly fewer calls in untagged or TGM2- inhibited conditions (Fig 3D). Setting stringent conditions, 66 modified sites were detected from 51 proteins (Table 1).

**Table 1.**
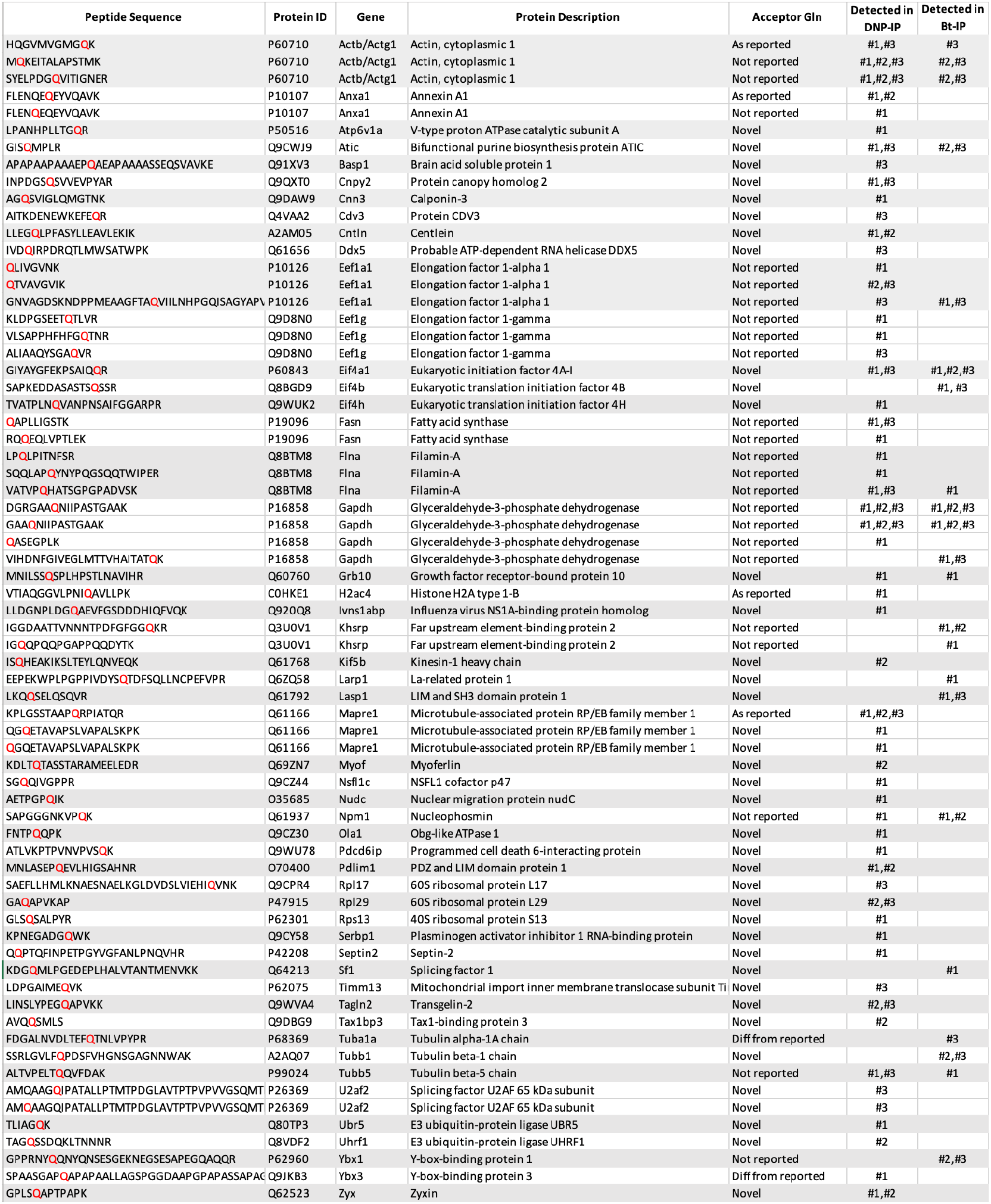
Combined list of polyaminated peptides identified with the different tagged probes. Acceptor glutamines shown in red. Last two columns indicate in which MS run the peptide was detected. Previously identified acceptor glutamines are referenced in (12) (19) (20). The Transdab Wiki (19) lists TGM2 polyaminated targets and their primary reference. “As reported” means the acceptor Gln has been previously reported and corresponds with our identification. “Not reported” refers to proteins that have been identified as polyaminated, but without identification of the acceptor Gln which are identified here for the first time. “Diff from reported” indicates that a different Gln was previously identified as polyaminated. “Novel” indicates a protein and its Gln acceptor site identified in this study without any prior report.

In vitro labelling of TGM2-KO lysate with Bt-PA did not yield any modified peptides (Fig 3D). All 6 detections came from the unlabelled control, of which 2 were immunoglobins used in the post-reaction pull-down. This suggests that these detections are spurious due to the higher sensitivity settings to detect any modified peptide present.

### TGM2 polyaminated targets

Identified peptides are curated in Table 1. While many peptides were detected in a single run, nevertheless many more were picked up multiple times. There was a significant overlap between the glutamine acceptor sites identified using the two different tags, lending confidence to the approach.

We found a substantial number of novel protein targets that have not been previously reported as TGM2 polyamination targets. We also identified hits that were previously reported as being polyaminated, but except for a few examples the glutamine acceptor was unidentified until now. The convention used to annotate the glutamine acceptor sites was as follows: (i) Novel, protein and acceptor Gln only detected in this study, (ii) Not reported, target has been reported to be polyaminated but Gln acceptor was not identified, (iii) As reported, target has been reported with polyamination at the same site, (iv) Different from reported, target has been reported but with a different Gln acceptor from this study.

Many of the targets were structural proteins (Actb/Actg1, Transgelin2, Pdlim1, Centlein, Mapre1, Zyxin, Filamin A, Annexin A1, Tubulin alpha-1a, Tubulin beta-1, Tubulin beta-5). The detected peptides are identical between Actb/Actg1 and we cannot distinguish which isoform, or if indeed both are modified. Targets in the translation machinery were also identified (Eef1a1, Eef1g, Eif4A1, Eif4b, Eif4h, Rps13, Rpl17, Rpl29). Interestingly, the modified glutamines (Q47, Q431) on Eef1a1 are on similar conserved motifs within the protein, suggesting some specificity in their selection. All these targets, including Gapdh and histone H2ac4 are abundant and could indicate an identification bias towards well represented proteins.

Among the other hits, polyamination at Q475 on Grb10 is of particular interest as it lies between S474 and S476. These serines are phosphorylated by the mTORC1 kinase and play a regulatory role in mTOR signalling (12, 15, 16). The juxtaposition of a potential polyaminated site between these 2 regulatory serines may indicate cross-talk between different modifications. The glycolytic enzyme Gapdh was modified on 3 sites with Q202 polyamination found in every run. Beyond its essential role in glucose catabolism, Gapdh has many unrelated moonlighting functions and has extensive PTMs (17). Interestingly, one of the polyaminated glutamines (Q183) lies between a threonine and lysine. These two sites are co-modified by phosphorylation and sumoylation respectively (18). As with Q475 on Grb10, the proximity of a polyaminated site with a regulatory hotspot may indicate an as yet undescribed role for polyamination in signaling cross-talk.

### Validation of in vivo Grb10 polyamination

As these targets were identified from in vitro polyaminated material, we wanted to validate if this is also the case in vivo. Grb10 was chosen as an example to validate in vivo polyamination, and FLAG-tagged Grb10 was over-expressed in CB1. Grb10 is expressed at low levels and the endogenous protein is unlikely to compete with the over-expressed construct for the polyamine label (Fig 4A). In vivo labelling was performed by Bt-PA incorporation into intact cells and FLAG-Grb10 protein was enriched from cell lysates by FLAG purification. FLAG-purified proteins were probed for the presence of a Bt-PA tag by immunoblot. As shown, Bt- polyaminated proteins were present in the FLAG-Grb10 pulldown (Fig 4B) indicating incorporation into FLAG-Grb10 by in vivo polyamination.

**Figure 4.**
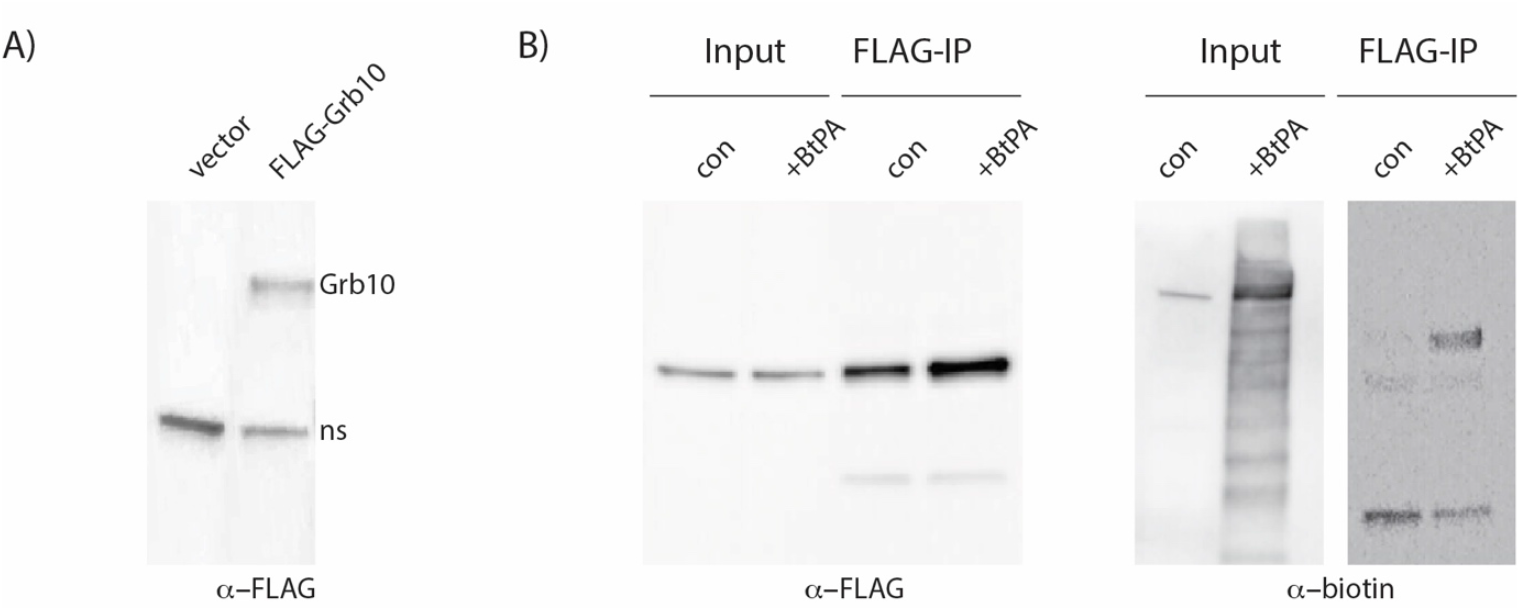
(A) Cloning and overexpression of FLAG-Grb10 in CB1. (B) Immunoblots showing FLAG IP of FLAG-Grb10 from control and Bt-PA treated CB1 cell lysate. The same samples were probed for the presence of the biotin tag using anti-biotin antibody.

## Discussion

Our study is the first proteomic survey of TGM2 polyamination since Yu et al. in 2015 (12). They identified 233 target proteins compared to 51 by us. There was only a small overlap of 7 proteins between both datasets (Filamin A, Histone H2ac4, Khsrp, Mapre1, Tuba1a, Tubb5, Ybx3). Nevertheless, in 2 of these cases, H2ac4 and Mapre1, the Gln-acceptors identified were identical which is highly unlikely to be by chance given the small sample size. The small overlap could be due to cell, tissue or species differences (mouse liver CB1 vs. human cervical HeLa), or methodology (their proteins were pulled down using an anti-spermine antibody). There was also an overlap with the Transdab database (19) that lists 156 TGM2 polyaminated targets. These hits were Actb/Actg1, Annexin A1, Eef1a1, Eef1g, Filamin A, Gapdh, and Nucleophosmin 1. Only a few glutamine acceptor sites are identified on Transdab but there was one common hit with the same Gln acceptor site (Annexin A1). We also found several hits in common with an earlier study by Orru et al. (Eef1a1, Eef1g, Fasn, Filamin A), however there were no Gln-acceptor sites identified in that study (20).

The remaining 36 hits were novel to this study. There was no apparent motif flanking the acceptor-Glns to indicate a specific recognition motif for TGM2 polyamination. A recent study on histone polyamination by TGM2 concluded that the main criteria dictating modification is steric accessibility of the polyaminated site (21). Our observation that polyamination of Gapdh and Grb10 are at modification hotspots appears to be in line with this conclusion.

We did not perform a gene ontology analysis of our hits to identify particular pathways or processes that may be over-represented. Our reluctance to do so was due to the small sample size. Additionally, as the overwhelming majority of pulled-down proteins were unable to be identified in the MS analysis, together with an apparent bias towards abundant proteins, any such analysis would be potentially misleading.

A limitation of this study is that the polyamine tags employed, Bt-PA and DNP-PA, have a 5-carbon long polyamine chain making them structurally similar to cadaverine instead of the more physiologically abundant and relevant polyamines (putrescine, spermidine and spermine). Unfortunately, tagged versions of putrescine, spermidine or spermine compatible with enrichment strategies are not commercially available. Other methods of labelling and enriching polyaminated proteins have been described such as click chemistry (22), and in vitro crosslinking (23). Having a variety of orthogonal approaches to enrich post-polyaminated proteins will lend confidence to identification if the same targets are identified via different routes.

Nevertheless, the main obstacle appears to be at the level of proteomic analysis and detection of polyaminated peptides. We observed that despite pulling down a large number of tagged proteins (Fig 3B, C), only very few targets were identified by MS, suggesting that there is substantial attrition of material during sample preparation or MS analysis. This is a major hurdle in the proteomic study of polyamination and remains an outstanding issue in the field.

## Acknowledgements

<u>M.N.H.</u> acknowledges support from the Swiss National Science Foundation (310030_192493)

## Legends

Table1. Combined list of polyaminated peptides identified with the different tagged probes. Acceptor glutamines shown in red. Last two columns indicate in which MS run the peptide was detected. Previously identified acceptor glutamines are referenced in superscripts (12) (19) (20). The Transdab Wiki (19) lists TGM2 polyaminated targets and their primary reference.

## Material and Methods

### Cell culture

CB1 cells are immortalized cells derived from a mouse liver cancer tumor (14). The cells are knocked out for Pten and Tsc1 and have constitutively active mTORC1 and mTORC2 activity. Cells were cultured in in Iscove’s medium supplemented with 10% fetal bovine serum, 2 mM L-glutamine, penicillin (100 U/ml), and streptomycin (100*μ*g/ml) at 37°C and 5% CO2. FLAG- Grb10 (Origene RC220715L3) and FLAG-GAPDH (Origene RC202309L3) were transfected into CB1 using JetPrime reagent and transfectants selected by puromycin selection (1*μ*g/ml). TGM2-KO cells were obtained by Crispr knockout (Santa Cruz sc-423375).

### FLAG-IP

Cell pellets were scraped in ice-cold PBS containing protease and phosphatase inhibitors. Pellets were dounced 30x in 2ml PBS and cleared by centrifugation at 20000g, 20min, 4°. 50*μ*l of washed anti-FLAG beads was added to 500*μ*g cleared lysate (at 1*μ*g/*μ*l concentration) and rotated for 4-6 hours at 4°. Beads were washed 3×1ml ice-cold PBS and bound proteins eluted with Flag peptide (2×30*μ*l, 0.5*μ*g/ml).

### Immunoblotting

Cells pellets were lysed in M-PER buffer and total protein (20 to 40 mg) resolved on SDS-PAGE gels. Proteins were blotted onto nitrocellulose membranes and probed with anti- DNP (SantaCruz sc-69698) or FLAG antibodies (Sigma Anti-FLAG M2 F1365). Biotin was probed with an HRP-Streptavidin conjugate (Sigma RABHRP3). TGM2 was probed with TGase2 Antibody (4G3), SantaCruz sc-73612).

### Polyamination reactions

<u>In vitro:</u> CB1 cell whole cell lysates were made using M-PER extraction buffer (ThermoFisher) containing protease inhibitors (cOmplete, Roche, 0.5mM PMSF) and phosphatase inhibitors (PhosSTOP, Roche). 100*μ*g of WCL lysate was diluted in 50mM MOPS, 5mM CaCl2 and 1mM Biotin-pentylamine (Bt-PA, Sigma) or DNP-pentylamine (DNP-PA, Zedira) was added. The reaction was incubated at 37° for 3 hours. 100*μ*M Z-DON was added for negative controls. At the end of the reaction, reaction products were passed through a size exclusion column (3kD cut-off) to remove unincorporated Bt-PA and DNP-PA.

<u>In vivo:</u> Confluent CB1 cells were directly treated with 1*μ*M ionomycin and 1mM labelled polyamines Bt-PA or DNP-PA. Cells were incubated for 4 hours before harvesting by scraping. Downstream processing of cell lysates was similar as for in vitro samples.

### Enrichment of polyaminated proteins

Post-reaction, cell lysates were diluted to 800*μ*l in 0.3% CHAPS buffer and anti-biotin (SantaCruz sc-101339AC) or anti-DNP (SantaCruz sc-69698AC) beads were added. Tubes were rotated for 4-6 hours at 4°. Beads were washed 3×1ml in ice-cold 0.3% CHAPS buffer and bound proteins either eluted by boiling for immunoblotting, or trypsin digested off beads for MS.

### Tryptic digest and LC MS/MS analysis

Affinity purified samples were subjected to on-bead digestion (adapted from PMID: 20479470). Resin was washed thrice with detergent free wash solution and collected by centrifugation. Proteins were eluted by incubation in 1.6 M Urea, 100 mM Ammonium bicarbonate, 5 *μ*g/ml trypsin, pH 8 for 30 min at 27°C shaking at 1200 rpm followed by two incubations in 1.6 M Urea, 100 mM Ammonium bicarbonate, 1 mM TCEP, pH 8. After each incubation, resin was collected by centrifugation and supernatants were collected and pooled. TCEP and Chloroacetamide were added at a final concentration of 10 mM and 15 mM, respectively, and samples were incubated for 1 h at 37°C shaking at 600 rpm prior to the addition of 0.5 *μ*g trypsin and incubation for 12 h at 37°C shaking at 300 rpm. Tryptic digest was acidified (pH<3) using TFA and desalted using C18 reversed phase spin columns (Microspin, Harvard Apparatus) according to the manufacturer’s instructions. Peptides were dried under vacuum and stored at -20°C.

Dried peptides were resuspended in 0.1% aqueous formic acid and subjected to LC–MS/MS analysis using a Q Exactive Plus Mass Spectrometer fitted with an EASY-nLC 1000 (both Thermo Fisher Scientific) and a custom-made column heater set to 60°C. Peptides were trapped on a PepMap Neo Trap Cartridge (Thermo Fisher Scientific) and resolved using a RP-HPLC column (75um × 30cm) packed in-house with C18 resin (ReproSil-Pur C18–AQ, 1.9 um resin; Dr. Maisch GmbH) at a flow rate of 0.2 ul/min. The following gradient was used for peptide separation: from 5% B to 10% B over 5 min to 35% B over 45 min to 50% B over 10 min to 95% B over 2 min followed by 18 min at 95% B. Buffer A was 0.1% formic acid in water and buffer B was 80% acetonitrile, 0.1% formic acid in water.

The mass spectrometer was operated in DDA mode with a total cycle time of approximately 1s. Each MS1 scan was followed by high-collision-dissociation (HCD) of the 10 most abundant precursor ions with dynamic exclusion set to 45 seconds. MS1 scans were acquired in Profile mode with a scan range from 350 – 1600 m/z, AGC target set to 3e6, a maximum injection time of 100 ms and a resolution of 70000 FWHM (at 200 m/z). MS2 scans were acquired in Centroid mode with a scan range from 200 – 2000 m/z, AGC target set to 1e5, a maximum injection time of 100 ms and a resolution of 35000 FWHM (at 200 m/z). Singly charged ions and ions with unassigned charge state were excluded from triggering MS2 events. The normalized collision energy was set to 27%, the mass isolation window was set to 1.4 m/z and one microscan was acquired for each spectrum.

The acquired raw-files were searched using MSFragger (v. 3.7) implemented in FragPipe (v. 19.1) against a Mus musculus database (consisting of 17085 protein sequences downloaded from Uniprot on 20220222) and 392 commonly observed contaminants. The default LFQ-MBR workflow was used. with minor modifications: In the MSFragger Spectral Processing tab, “Require precursor” was unchecked, MSBooster was disabled, in the Quant (MS1) tab “MBR top runs” was set to 100000, “Top N ions” was set to 3 and “Min freq” was set to 0.5. A variable modification on Q with a delta mass of either 251.0906 Da (DNP-PA) or 311.16675 Da (Bt-PA) was added.

## Notes

### Competing Interest Statement

The authors have declared no competing interest.

